# Ultra-parallel ChIP-seq by barcoding of intact nuclei

**DOI:** 10.1101/276469

**Authors:** L. Arrigoni, H. Al-Hasani, F. Ramírez, I. Panzeri, D.P Ryan, D. Santacruz, N. Kress, A. Pospisilik, U. Böenisch, T. Manke

**Author notes:** Correspondence should be addressed to Thomas Manke or Laura Arrigoni.

## Abstract

Chromatin immunoprecipitation followed by deep sequencing (ChIP-seq) is an invaluable tool for mapping chromatin-associated proteins. However, sample preparation is still a largely individual and labor-intensive process that hinders assay throughput and comparability. Here, we present a novel method for ultra-parallelized high-throughput ChIP-seq that addresses the aforementioned problems. The method, called RELACS (Restriction Enzyme-based Labeling of Chromatin *in Situ*), employs barcoding of chromatin within intact nuclei extracted from different sources (e.g. tissues, treatments, time points). Barcoded nuclei are pooled and processed within the same ChIP, for maximal comparability and significant workload reduction. The choice of user-friendly, straightforward, enzymatic steps for chromatin fragmentation and barcoding makes RELACS particularly suitable for implementation large-scale clinical studies and scarce samples. RELACS can generate ChIP-seq libraries from hundreds of samples within three days and with less than 1000 cells per sample.

---

INTRODUCTION

Chromatin immunoprecipitation followed by deep sequencing (ChIP-seq) is a key technique for exploring the genomic location of bound proteins and histone modifications, which has significantly contributed to our understanding of gene regulation and epigenetic changes in healthy and diseased cells. Despite many protocol improvements 1–4, ChIP-seq still suffers from limitations imposed by the complexity of the protocol: limited throughput, low efficiency for proteins weakly bound to chromatin, and a large amount of cellular material needed. The labor-intensive nature of ChIP-seq protocols impede parallel processing of multiple samples, eventually hindering comparability and standardization of large-scale studies and clinical applications with scarce material. In conventional ChIP-seq, chromatin extraction, fragmentation (using sonication or micrococcal nuclease) and enrichment by immunoprecipitation are carried out for each individual sample. After DNA purification, library preparation protocols are applied prior to multiplexed sequencing. Recent ChIP-seq methods 1,2,5 aim to increase throughput by employing molecular barcoding, which facilitates downstream parallel processing of multiple samples within the same ChIP. Barcode sequences are ligated to each starting sample and are used to identify signals from each cell population during data analysis.

Though improved, these methods are still limited. For example, some high-throughput procedures are applicable only to histone modifications using fresh unfixed material 2, or involve repeated ChIP applications (re-ChIP) that severely limits assay throughput and does not significantly reduce the workload compared to standard ChIP-seq ^1^. Barcoding of sonicated chromatin, as used in iChIP and derived techniques ^1^,^6^, requires capturing chromatin with beads to remove chromatin from a detergent-containing solution, which would inhibit barcode ligation barcodes. Other methods ^2^,^5^ do not require re-ChIP, but rely on micrococcal nuclease (MNase) digestion of chromatin. This is prone to chromatin over-fragmentation because of exonuclease activity, especially when dealing with variable amounts of cells^7^,^8^. Therefore, MNase is far from optimal when investigating transcription factors or histone modifications associated with open chromatin or when using limited material

Here, we introduce a high-throughput user-friendly ChIP-seq method based on the enzymatic cutting and barcoding of chromatin inside intact nuclei. Once multiple cell populations are barcoded, they are pooled for parallelized ChIP-seq. The method, called RELACS (Restriction Enzyme-based LAbelling of Chromatin *in Situ*), addresses the shortcomings of current multiplexed ChIP-seq techniques: it can be used for a broad spectrum of epitopes and cell types, it is robust to variation of cell numbers or experimental conditions, and its throughput improves comparability while decreasing experimenter workload. As with other barcoding strategies, this approach can also be used for quantitative ChIP-seq analyses.

## RESULTS

### Method description

The protocol starts by isolating clean nuclei from fixed cells, which enhances protocol standardization across cell types ^9^ (**Fig. 1**). Next, restriction endonucleases with a high number of cut sites are used to create short chromatin fragments, while keeping the nuclear envelope intact. Digestion is performed at room temperature to reduce temperature-induced crosslink reversal ^10^. Chromatin is kept inside permeabilized nuclei throughout the entire process of digestion and barcoding. Maintaining chromatin in nuclei facilitates downstream wash and pooling steps without increasing sample volumes (since one can simply precipitate the nuclei). Inside the intact nuclei, customized adapters containing nuclear barcodes are ligated to both ends of the chromatin fragments (**Supplementary Fig. 1**). This conventional adapter ligation strategy is both robust to variation of cell number and practical to implement in various laboratory and clinical settings. Lastly, chromatin containing different barcoded samples can be aliquoted prior to ChIP for the investigation of multiple epitopes. A second barcode is inserted via a PCR step to mark different ChIP experiments or controls. In addition to a simplified workflow and enhanced comparability, the multiplexing low cell number samples also allows easier handling and reduces potential loss of chromatin during immunoprecipitation.

**Figure 1:**
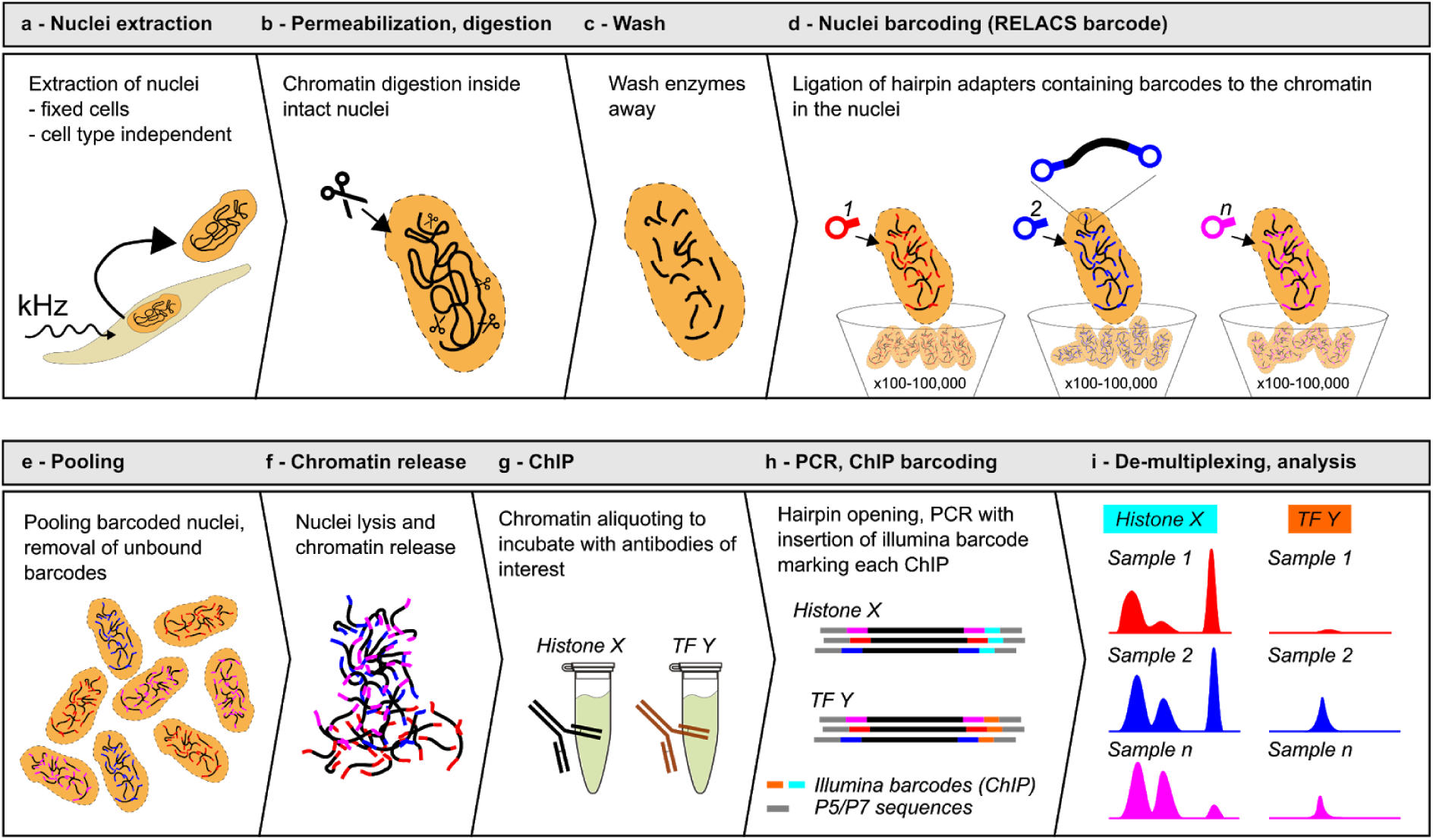
RELACS workflow. Overview of the RELACS method. The protocol facilitates barcoding multiple cell populations, which can be pooled and investigated for multiple epitopes within the same run. The method starts by isolating nuclei from a pool of formaldehyde-fixed cells, using sonication to reduce cell type dependency 9 (a). The nuclear membrane is permeabilized to allow entrance of enzymes, followed by DNA barcodes. One or more restriction endonucleases with a high frequency of recognition sites are used to fragment chromatin (b). Nuclei are washed to remove active restriction enzymes (c). Hairpin adapters harboring barcodes are ligated to both the ends of the fragmented chromatin inside the nuclei. The barcoding has been tested using 100 to 100,000 nuclei without the need to change protocol conditions (d). Cell populations marked with specific barcodes are pooled (e), concentrated and lysed to release chromatin into solution (f). Chromatin is split and incubated with the antibodies of interest (g). After ChIP washes and DNA purification (not illustrated), only DNA that harbors nuclei barcodes at both ends is PCR amplified to complete library construction. PCR amplification appends an Illumina barcode onto mark each fragment (h). Sequenced libraries are de-multiplexed by Illumina barcode, to retrieve ChIP information, and then nuclear barcode, to identify the initial cell population (i). The RELACS protocol is very fast and ChIP-seq libraries can be generated for hundreds of samples within three days. For a more detailed description of library construction (nuclei barcoding and PCR amplification) see **Supplementary Fig. 1**.

### RELACS results are comparable to traditional ChIP-seq

We validated RELACS using a well-studied human cell line (HepG2) for which many external references are available. Here, we generated six histone modification maps for IHEC class I epigenomes (H3K27ac, H3K4me3, H3K4me1, H3K36me3, H3K9me3, H3K27me3), the insulator protein CTCF, as well as co-factor p300 using a traditional ChIP-seq protocol as well as RELACS. All marks show very high quality results, as illustrated in **Figure 2a** and quantified by quality metrics such as mapping rates, duplication rates and fraction of reads in peaks (FRiP) scores. For FRiP score calculation, we used annotated peak regions from the ENCODE consortium (see Methods). Genome-wide coverage tracks and average profiles around known target regions show excellent agreement of the RELACS method with a traditional approach (**Fig. 2a, b**). While the traditional approach used 1x 100000 cells per ChIP, RELACS data was generated from 20 technical replicates (20 barcodes x 5000 cells per ChIP) and merged for subsequent analysis. Barcode distribution is consistent across epitopes (**Supplementary Fig. 2a**) and there is no apparent cross-barcode contamination (**Supplementary Fig. 2b**). The overall mapping rates are very high (>99%) and we observed only small fractions (3-5%) of disconcordantly and multi-mapping reads (**Supplementary Data 1**, **Supplementary Fig. 2c**). These reads were excluded from subsequent analyses.

**Figure 2:**
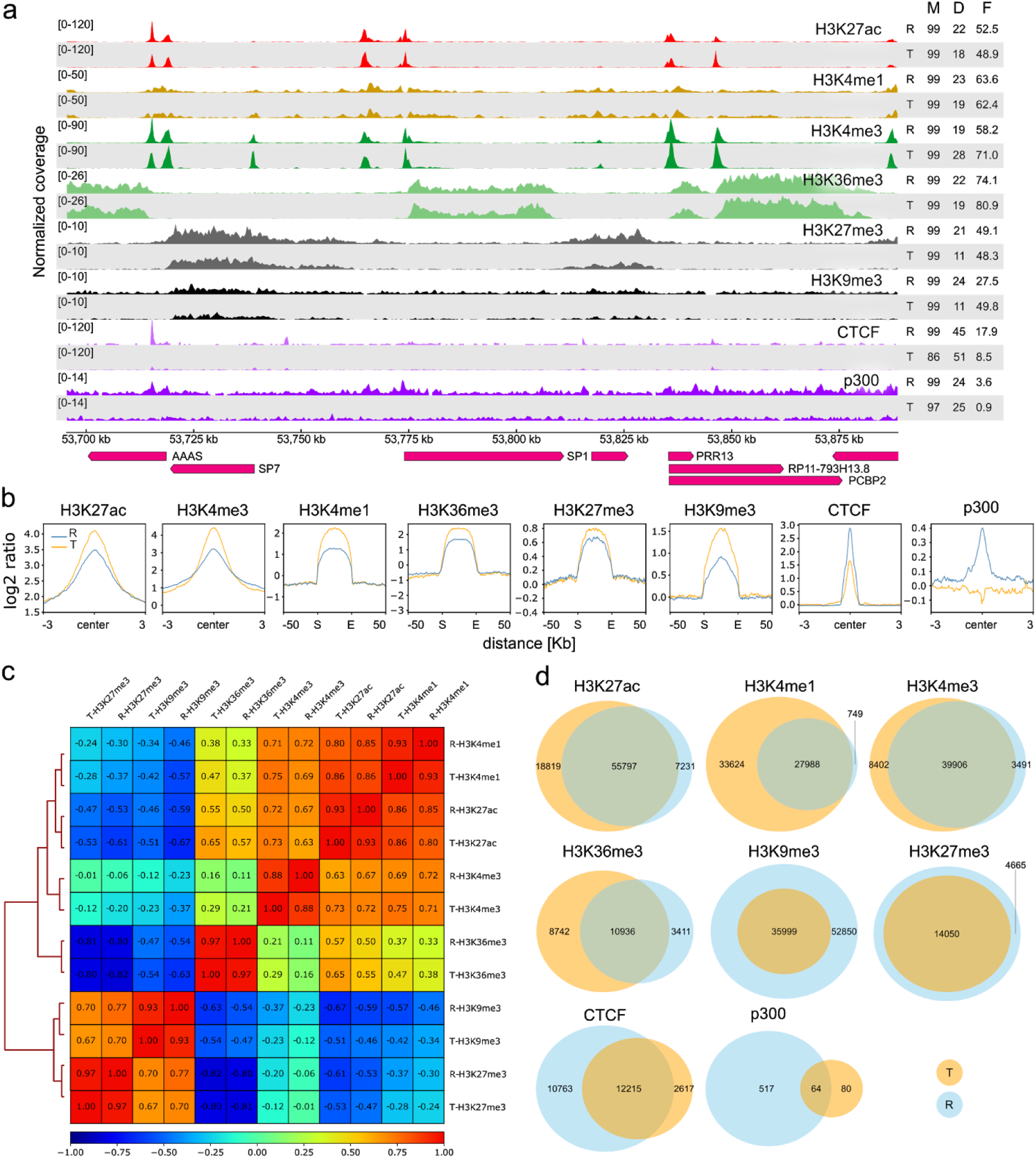
RELACS Validation. Using the HepG2 cell line as a model system, we compare results from RELACS with those from a traditional ChIP-seq method. (**a**) Data tracks of 6 histone marks (H3K4me1, H3K27ac, H3K4me3, H3K36me3, H3K27me3, H3K9me3), transcription factor (CTCF) and a co-factor (P300) are shown. There is a high correlation within marks prepared with RELACS, designated with an R, and a traditional method, designated with a T9. On the side, mapping rate (M), duplication rate (D) and the fraction of reads in peaks (FRiP, F) are assigned to each track. (**b**) Signal scores of RELACS vs. traditional ChIP over peaks from ENCODE (see online methods). For sharp marks we centered the profile around the annotated peak center and added 3 kb to either side. For broad marks we scaled the regions to 50 kb and added a flanking region of 50 kb. (**c**) Pearson correlation of log2-ratios (ChIP/Input) for RELACS (R) and Traditional (T) samples. The correlation was computed for 10 kb bins. **(d)** Venn diagrams for overlaps of enriched regions called by MACS2 (for sharp marks: H3K27ac, H3K4me3, CTCF, p300) and histoneHMM (for broad marks: H3K4me1, H3K36me3, H3K9me3, H3K27me3).

RELACS data also shows proper signal distributions of transcription factors and co-factors such as CTCF and p300, for which the traditional approach is much poorer or fails when using small cell amounts (**Fig. 2b**). Such success cannot be claimed by other multiplexed ChIP-seq protocols ^2^, which cannot be applied to transcription factors and also reportedly have problems in mapping sharp histone marks. The resolution achieved with restriction enzymes is comparable to sonication-based fragmentation (**Fig. 2b**), indicating that our choice of restriction enzyme results in data faithfully conveying the underlying signal distributions.

A more detailed analysis of restriction site occurrence in the genome (cutability) and restriction site preference (accessibility) indicates that both are reduced in heterochromatic regions (**Supplementary Fig. 3**). Other repressed regions are comparable between RELACS and the traditional protocol, while open chromatin regions show higher accessibility for RELACS. As with all ChIP-seq studies, we account for such biases by employing an input control for normalization and peak calling. The genome-wide correlations between the log2-ratios (ChIP/Input) from the traditional method and RELACS are high (0.88-0.97, **Fig. 2c**) and we observe large overlaps between peaks (**Fig. 2d**). This also suggests that standard peak calling algorithms (such as MACS2 ^11^) can be directly used to analyse RELACS data. The RELACS fragmentation strategy does not appreciably affect genomic coverage, as indicated by the comparable traditional method results (**Supplementary Fig. 4**). The only exception to this is in heterochromatic regions, where coverage in RELACS is lower at the same sequencing depth. Importantly, regions of prime experimental interest are better covered at lower depths.

### RELACS works reliably with low cell numbers

Barcoding intact nuclei before any further treatment facilitates pooling of scarce samples prior to ChIP, as previously shown^2^,^12^. Such pooling results in sufficient material for immunoprecipitation and quality controlling, which is typically only possible given samples with abundant material. Here we tested the sensitivity limits of RELACS by barcoding seven replicates with 10000, 1000, or 100 HepG2 cells each and compared the ChIP-profiles around known target regions. We found that RELACS can produce robust and reproducible profiles down to 100 cells for histone marks and 1000 cells for transcription factors, beyond which the signal-to-noise ratio deteriorates **(Fig. 3)**. To quantify signal reduction with decreasing cell numbers, we computed the fraction of reads that overlap ENCODE annotated peak regions and found that it ranges from from 60% for 100000 cells to 20% for 100 cells (**Supplementary Fig. 5a**). Simple peak calling with default parameters generates consistent regions between the traditional protocol with 100000 cells and the RELACS protocol with only 1000 cells (**Supplementary Fig. 5b**).

**Figure 3:**
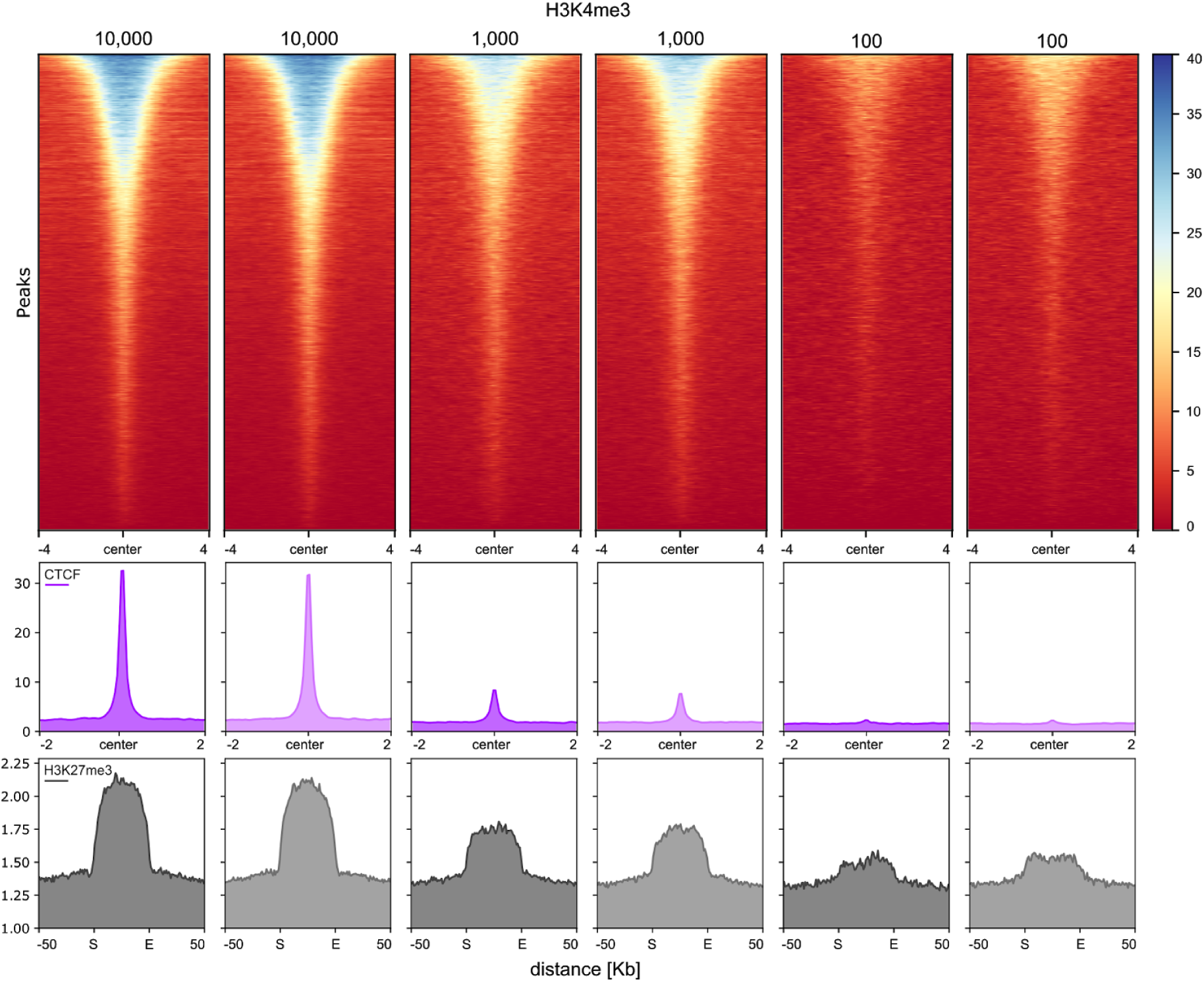
RELACS sensitivity using low cell numbers. Comparison of RELACS results using 10,000, 1000 and 100 HepG2 cells as starting material. For each cell number, 7 technical replicates were barcoded and pooled for ChIP-seq and computationally demultiplexed before analysis. This figure shows two replicates for each cell number. The top heatmap shows the 1x-normalized read coverage for the H3K4me3 histone mark at its respective ENCODE peaks. The coverage is shown centered at the peak with 4 kb flanks on each side. Similarly, for CTCF (middle panel) the normalized coverage at the peak center shown together with 2 kb flanking regions. For H3K27me3 (last panel) normalized coverage at each ENCODE peak is scaled to 50 kb (S: start position; E: end position) and flanking 50 kb are shown.

### RELACS parallelizes over tissues

We applied RELACS to barcode nuclei from 8 different mouse tissues and two biological replicates for a total of 16 x 25000 cells per ChIP. For each histone mark (H3K36me3, H3K27ac, and H3K27me3) only a single ChIP on pooled-and-barcoded nuclei was performed for maximal simplicity and comparability (**Fig. 4a**). After sequencing, the pooled samples were demultiplexed according to the nuclear barcodes to assign all reads to their tissues of origin. We observed high-quality enrichment profiles (**Fig. 4b**) and mapping rates, duplication rates and the FRiP scores (**Supplementary Data 2**). Our results are highly reproducible, as seen by clear replicate clustering in PCA plots for all analyzed histone marks (**Fig. 4c**). A control sample (“input”) is prepared for each ChIP experiment, which not only accounts for the usual biases in the protocol, but also faithfully conveys the differences in initial sample amount and barcoding efficiency. ChIP signals of each multiplexed sample can, therefore, be normalized against their respective input controls and signals between samples can be quantitatively compared. Using multiplexed H3K36me3 ChIPs and inputs on identical HepG2 samples, we show that we can reproduce the relative scaling factor (which should be 1) within 10% (**Supplementary Fig. 6a**). This same procedure, then, can be applied to multiplexed tissues as well, to allow quantitative comparisons between them (**Supplementary Fig. 6b**). In some cases (e.g. brain), we observe consistently high scaling factors, indicating that these tissues may have overall higher levels of H3K36me3 compared to others.

**Figure 4:**
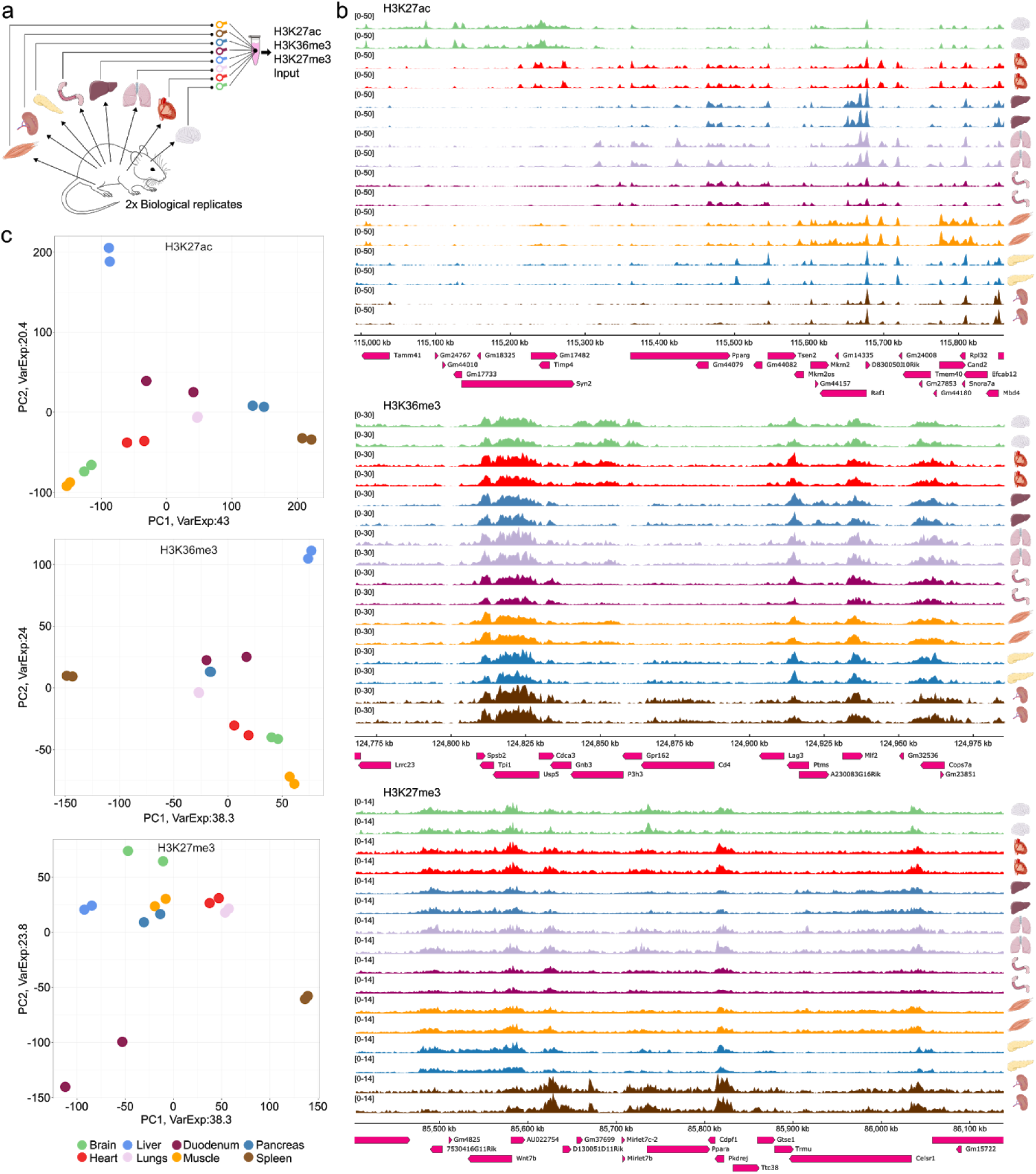
Analysis of multiple tissues and biological replicates in a single ChIP. (**a**) Schematic representation of the high-throughput RELACS ChIP-seq experiment. Eight mouse organs (brain, heart, lungs, liver, duodenum, pancreas, spleen, skeletal muscle) were extracted from two wild-type mice. Organs were independently homogenized, fixed, and nuclei were extracted. After chromatin digestion inside intact nuclei, different hairpin nuclear barcodes were used to mark nuclei extracted from each organ. After barcoding nuclei were pooled and lysed to release chromatin. Chromatin was split to investigate three histone modifications (H3K27ac, H3K36me3, H3K27me3) plus input for normalization control. (**b**) Tracks of three different histone marks (H3H27ac, H3K36me3, H3K27me3) with 8 different mouse tissues derived from two biological replicates are shown. (**c**) Principal component analysis illustrates a clear and consistent separation of tissues for all three histone marks where more than 60% of variation is explained (“VarExp” in the figure) by the first two components. Cases where only one replicate is visible indicate perfect overlap within the limits of resolution.

## DISCUSSION

We have introduced an ultra-parallel and highly standardized approach to the generation and analysis of ChIP-seq data. It can be established easily in any molecular biology lab without specialized equipment. Compared to MNase-based protocols (e.g. Mint-ChIP^2^), RELACS is simpler, as it does not need strict control of MNase concentrations to prevent over-cutting open chromatin regions and to ensure result consistency^13^. Unlike Mint-ChIP, it can also be applied to fixed samples, allowing profiling of transcription factors and co-factors easily lost by native ChIP approaches. The ability to label and pool large number of samples opens up the possibility to conduct controlled ChIP-seq experiments with very small cell numbers. We have obtained reliable and reproducible histone profiles for as few as 100 cells and transcription factor profiles from 1000 cells. We do not exclude the possibility that ChIP using even lower cell numbers can be performed, by merging a higher number of barcoded small cell populations, or by adding un-barcoded carrier chromatin prior to immunoprecipitation (to reduce the noise associated with low input samples)^14^,^15^.

Compared to the iChIP protocol^1^ for small cell numbers, RELACS uses intact nuclei throughout chromatin fragmentation and barcoding. This eliminates the need for an additional ChIP step just to purify chromatin prior to barcode ligation. As a result, RELACS provides a simple and highly standardized workflow for all cell types and a wide range of sample and cell numbers.

We have carefully investigated the potential limitations and biases due to the restriction site based RELACS methodology. Overall, the genome coverage at a given sequencing depth is comparable to a traditional approach, with slightly reduced coverage of heterochromatic region and repetitive regions (due to reduced cutability and accessibility) and concomitantly increased coverage of many types of open regions. As with all other ChIP-seq protocols, we also observe a bias towards open chromatin^16^, where most of the studied marks are located (active histone marks, CTCF, p300). These biases are accounted for by using input controls, which is already done in standard algorithms for peak calling. For transcription factors (CTCF) the RELACS protocol is more sensitive, and for co-factor p300 only the RELACS method was able to obtain a signal in known target regions. Importantly, for sharp marks we achieved the same resolution as with sonication based methods. In our analyses with extremely small cell numbers (<1000 cells) and sharp marks (CTCF), we observed a clear reduction in library complexity, which results in high duplication rates at a given sequencing depth. In this work we have aggressively filtered and removed all duplicate alignments, because it is currently not possible to distinguish technical PCR duplicates from biological duplicates derived by restriction digestion. Future improvements may employ unique molecular identifiers to resolve this issue.

Given the broad applicability of RELACS (universality across tissues, cell numbers, histone or transcription factor ChIP-seq), we propose that it can be used as a reference protocol for ultra-parallel handling of limited clinical samples and large-scale comparative studies. Though we have applied RELACS to multiple mouse tissues, the protocol is compatible with more stringent purification techniques (e.g. nuclear sorting), allowing the production of epigenetic profiles from any combination of samples. Moreover, the protocol could also be integrated into other assays, such as DNA methylation analysis or chromatin conformation studies.

## ACCESSION CODES

Data has been uploaded to the Gene Expression Omnibus (GEO) with accession code: GSE111000.

## ACKNOWLEDGEMENTS

We would like to thank Vivek Bhardwaj for help with oligo design and Ken Lam for providing cultured S2 cells. The results upon which this publication is based were funded by the Federal Ministry of Education and Research (BMBF) under the project number 01KU1216G (German Epigenome Programme - DEEP) and the CRC992 of the German Science Foundation (SFB “Medical Epigenetics”).

## AUTHOR CONTRIBUTIONS

L.A. performed and conceived the study, H.A., F.R. and T.M. performed data analysis and prepared figures, I.P. and A.P. supported the preparation of mouse experiments, D.P.R. aided the analysis and established the RELACS de-multiplexing pipeline, D.S. and N.K. helped with sequencing, U.B. provided experimental support and advice, T.M. supervised the project. L.A., F.R and T.M. wrote the manuscript.

## METHODS

### Cell culture

HepG2 liver hepatocellular carcinoma (ATCC, HB-8065TM) were cultured in Eagle‘s Minimal Essential Medium (EMEM, Lonza, 06-174) supplemented with 10% Fetal Bovine Serum (Sigma), 2 mM L-Glutamine (Lonza), 1.8 mM CaCl_2_, 1mM sodium pyruvate (Lonza) and penicillin-streptomycin mixture (100 units/ml, Lonza), at 37 °C at 5% CO_2_ in 10 cm plates, up to 80% confluency. S2 cells were cultured in Express Five SFM (Thermo Fisher Scientific) supplemented with glutamax, at 27°C.

### Mouse tissues

Experiments were performed on 9 week-old wild-type (WT) FVB/NJ littermate male mice. Mice were euthanized using carbon dioxide (CO_2_) before organ extraction. Dissected organs have been placed in room temperature D-MEM prior to fixation. All animal studies were performed with the approval of the local authority (Regierungspräsidium Freiburg, Germany)

### Cell fixation for cell line and tissues

Adherent HepG2 and S2 cells were fixed in 1% methanol-free formaldehyde (Thermo Scientific, 28906) in D-MEM (for HepG2 cells) or Express Five SFM (for S2 cells) for 15 minutes at room temperature under gentle shaking. Formaldehyde was quenched for 5 min by adding 125 mM glycine final concentration. Cells were rinsed twice with ice-cold PBS, harvested by scraping and pelleted (300 g, 10 min, 4 °C). For tissues, a small portion of each organ was finely chopped. Organ fragments were transferred into a Dounce homogenizer (loose pestle), covered with 1 ml of 1% formaldehyde in D-MEM, and dissociated with 3-4 strokes, followed by 15 minute incubation at room temperature. During incubation, the tissue suspension was filtered using a nylon 70 µM cell strainer (Falcon, 352350) to remove larger debris. 125 mM glycine final was added to the sample and the suspension was pelleted for 5 min at 500 g. Cell pellets were washed twice in PBS supplemented with a protease inhibitor cocktail (Roche, 11873580001) and aliquoted. Fixed cell pellets were stored at -80 °C until further usage.

### Chromatin preparation (for traditional sonication-based protocol)

Chromatin from HepG2 fixed cell pellets was prepared with the standardized method used for DEEP/IHEC consortia epigenome production, as previously described^9^. Briefly, nuclei were extracted using sonication (NEXSON) under these sonicator parameters: 75W peak power, 2% duty factor, 200 cycles/burst, for 90 seconds of treatment (Covaris E220 and 1 ml tubes, cat. No. 520130). Nuclei were pelleted, followed by sonication in shearing buffer (10 mM Tris-HCl pH 8, 0.1% SDS, 1 mM EDTA) using these sonicator parameters: 140W peak power, 5% duty factor, 200 cycles/burst, 15 minutes of treatment, 1 ml tubes. A chromatin aliquot was de-crosslinked, purified, and quality controlled for DNA concentration (Qubit DNA HS, Invitrogen, Q32851) and size distribution using capillary electrophoresis (Fragment Analyzer, NGS 1-6000 hs DNA kit).

### Nuclear barcodes construction

The sequence of hairpin barcode adapters is reported in **Supplementary Table 1.** Ultramer oligonucleotides were purchased from IDT (Integrated DNA Technology). Lyophilized oligos were resuspended in annealing buffer (10 mM Tris-HCl pH 8, 50 mM NaCl, 1 mM EDTA) to a final 100 mM concentration. Prior to hairpin formation, oligos were diluted to 15 mM in annealing buffer. For annealing, diluted oligos were heated at 95°C for 2 min in a thermoblock and afterwards switched off and slowly cooled down to room temperature.

### RELACS barcoded chromatin preparation

RELACS involves these key steps: nuclei extraction using sonication, nuclei swelling, nuclei digestion, nuclei wash, nuclei barcoding, nuclei pooling and chromatin release.

### 1. Nuclei extraction by sonication (NEXSON)

Formaldehyde-fixed pellets were resuspended in 1 ml of lysis buffer (10 mM Tris-HCl pH 8, 10 mM NaCl, 0.2% Igepal) supplemented with 1X protease inhibitor cocktail (Roche, 11873580001). Nuclei were extracted by sonication using the NEXSON approach^9^. For all cell types, parameters for nuclei extraction were the following: 75W peak power, 2% duty factor, 200 cycles/burst, 1 ml tubes (cat. No. 520130) using the Covaris E200 sonicator. Treatment time was stopped when nuclei extraction was satisfactory (over 70% of isolated nuclei). The following treatment times was used: 60-120 seconds (HepG2), 30 seconds (S2 cells), 45 seconds (liver, brain, spleen, lungs, skeletal muscle, duodenum), 30 seconds (pancreas, heart). After nuclei extraction, a small aliquot of nuclei was set aside to estimate DNA concentration and cell number from tissue samples: for this purpose, nuclei aliquots were shortly sonicated, de-crosslinked and DNA was purified using columns.

### 2. Nuclei swelling

Approximately 500,000 nuclei were pelleted (1000 g, 5 min), resuspended in 50 µl of 0.5% SDS and incubated at room temperature for 10 min. SDS was quenched adding 25 µl of 10% Triton X-100 and 145 µl of water. 25 µl of restriction enzyme buffer (CutSmart, NEB B7204S) and 2.5 µl of 100X protease inhibitor cocktail was added prior to enzyme incubation.

### 3. Nuclei digestion

Five units of the restriction enzyme CviKI-1 (NEB, R0710S) were added to every 100,000 nuclei. Samples were digested for 16 hours at 20 °C in a thermomixer under shaking (800 rpm). Note that the restriction endonucleases (used in RELACS) lack exonuclease activity and are easier to titrate upon variation of the input material. No significant differences are observed using 20 units of enzyme to digest approximately 10000 to one million fixed cells.

### 4. Nuclei wash

Restriction enzymes were removed by pelletting the nuclei (1000 g, 5 min) and then washing in 200 µl of nuclei wash solution (10 mM Tris-HCl pH 8, 0.25% Triton X-100, 0.1 mg/ml BSA). At this stage 10-20 µl of nuclei were set aside for digestion quality control. Remaining nuclei were pelleted down and resuspended in 10 mM Tris-HCl pH 8 (25 µl per 100 to 100,000 nuclei) for barcoding.

### 5. Nuclei barcoding

Digested chromatin inside nuclei was end-repaired and A-tailed using end-repair and A-tailing components from NEBNext Ultra II DNA library preparation kit (E7645L, NEB) with the following modifications, to enhance efficiency and to reduce handling volumes: 1.5 µl of end prep enzyme mix and 3.5 µl of reaction buffer were added to each 25 µl digested nuclei aliquot. Samples were mixed and incubated at 20 °C for 30 min followed by heat inactivation at 65 °C for 5 min. 1.2 µl of hairpin adapter nuclear barcodes were added to each sample. Ligation of barcodes to A-tailed chromatin was performed using components from NEBNext Ultra II DNA library preparation kit, with the following modifications: per sample 15 µl of ligation master mix and 0.5 µl of ligation enhancer were added, followed by 15 min incubation at 30 °C and 15 min incubation at 20°C. Barcoding efficiency is consistent between 100 to 100,000 nuclei using an unmodified protocol.

### 6. Nuclei pooling and chromatin release

Ligase was inhibited by adding 4.7 µl of 3 M NaCl into each well. All barcoded nuclei were pooled and pelleted down (5000 g, 10 min, 20 °C). Supernatant was removed and re-centrifuged at higher speed to increase recovery (11000 g, 5 min, 20 °C). Nuclei pellets were combined and re-suspended in shearing buffer (10 mM Tris-HCl pH 8, 0.1% SDS, 1 mM EDTA; approx. 130 µl per up to 500,000 nuclei maximum). Chromatin was released by sonication-assisted nuclei lysis by treating the samples for 5 minutes using these parameters: peak power 105 W, 2% duty factor, 200 cycles/burst, Covaris microtubes (520052), Covaris E220 sonicator. This barcoded chromatin was afterward used for ChIP.

### Digestion quality control

ChIP elution buffer (10 mM Tris-HCl pH 8, 1 mM EDTA, 1% SDS) was added to each nuclei aliquot to a final 100 µl volume. Samples werede-crosslinked and deproteinized by adding 4 µl of 5M NaCl, 2 µl 10 mg/ml DNase-free RNase A and 2 µl of 20 mg/ml proteinase K and incubating 30 min at 37 °C followed by 65 °C incubation for a minimum of 2 hours. Samples were purified using Qiagen PCR purification kit columns. DNA was quantified using a Qubit DNA HS kit and fragment size distribution monitored by capillary electrophoresis. Cell number for tissue samples was estimated from the total DNA amount, taking into account that each mouse diploid cell contains approximately 6.6 pg of DNA.

### Antibodies

The following antibodies targeting histone modifications were used for ChIP-seq (1 μg per ChIP for cell numbers below 100,000, 2 μg per ChIP for cell total cell numbers per ChIP below 250,000): anti-H3K27ac (C15410196, lot A1723-041D), anti-H3K4me3 (C15410003, lot A5051-001P), anti-H3K4me1 (C15410194, lot A1863-001D), anti-H3K36me3 (C15410192, lot A1847-001P), anti-H3K9me3 (C15410193, A1671-001P), anti-H3K27me3 (C15410195, lot A1811-001P), all from Diagenode. Transcription factors antibodies: CTCF (3 μg/ChIP, Abcam, ab70303), p300 (10 μg/ChIP, Santa Cruz, sc-585).

### ChIP

RELACS or sonication-based chromatins were diluted 1:2 in 1X buffer iC1 from iDeal ChIP-seq kit for histones (Diagenode, C01010173) and supplemented with 1X protease inhibitor cocktail (Roche complete EDTA-free, 11873580001). Diluted chromatin was divided into 200 μl aliquots and the antibody of interest has been added. For transcription factor ChIP, an additional 2.6 µl of 5M NaCl was added to each ChIP to adjust the final salt concentration to 140 mM. Automatic ChIP was performed using the SX-8G Compact IP-Star platform (Diagenode), immunoprecipitation buffers from the iDeal ChIP-seq kit for histones and A-conjugated magnetic beads (Diagenode, C03010020). The following IP-Star pre-programmed parameters were used for automated ChIP: “indirect ChIP method” using “200 µl ChIP volume”. Immunoprecipitation was carried out by incubating chromatin with the antibody for 10 hours at 4 °C. Immunocomplexes were captured by protein-A conjugated magnetic beads (3 hours of beads incubation at 4 °C) and washed four times using iDeal 1-4 wash buffers (5 minutes of incubation at each wash). After ChIP, eluates were recovered manually, RNase A-treated, de-crosslinked and deproteinized for 30 minutes at 37 °C and 4 hours at 65 °C. DNA was purified using Qiagen MinElute columns (Qiagen, 28006) with a final elution in 20 µl.

### Library amplification for RELACS ChIP-seq samples

18 μl of RELACS ChIP DNA was mixed together with 25 μl of PCR master mix from NEBNext Ultra II DNA library preparation (E7645L, NEB), 3 μl of USER enzyme and 2 μl of 10 μM universal PCR primer. 2 μl of 10 μM indexed primers were added to each respective ChIP. Hairpin adapters were opened by a 15 min incubation at 37 °C. 12 PCR cycles were used for enrichment of ligated fragments using the following program: hot start 98 °C 30 sec. Amplification: 98 °C 10 sec, 65 °C 75 sec. and a final extension at 65 °C for 5 min. PCR-amplified samples were purified twice using Ampure XP beads (Beckman Coulter) at 0.9 and 1X ratio.

### Traditional ChIP-seq library preparation

For each library, 1-5 nanograms of purified ChIP DNA was used. Libraries were prepared using the NEBNext Ultra II DNA Library Prep kit for Illumina (E7645L, NEB) with the following modifications. After adapter ligation, size selection was omitted and adapter-ligated fragments were purified using AMPure XP beads (Beckman Coulter) at 1X ratio. USER enzyme treatment was performed together with PCR enrichment using the following program: 15 min incubation at 37 °C, hot start 98 °C for 30 sec., then 12 cycles of 98 °C for 10 sec and 65 °C for 75 sec. A final extension was done at 65 °C for 5 min. Amplified libraries were purified twice using 0.9 and then 1X Ampure XP beads.

### Library quality control and sequencing

All libraries were quality-controlled for DNA concentration (Qubit DNA HS, Invitrogen, Q32851) and size distribution (capillary electrophoresis Fragment Analyzer, NGS 1-6000 bp hs DNA kit). Adapter-cleaned libraries were normalized to the desired molarity and then pooled accordingly with a 10% PhiX spike-in. Library pools were denatured and prepared for instrument loading following Illumina guidelines. Samples were sequenced paired-end with a read length of 75 bp on an Illumina HiSeq 3000 instrument.

### Demultiplexing, Mapping and Quality control

Using an in house written script the sequences were demultiplexed. Sequences without barcode or having more than one mismatch between barcodes on each paired-end mate were discarded from further analysis. HepG2 reads were mapped to the hs37d5 reference human genome and mouse reads were mapped to mm10 reference genome. snakePipes DNA-mapping workflow v0.3.2.1 (https://zenodo.org/badge/latestdoi/54579435) was used to compute quality controls using FastQC (https://www.bioinformatics.babraham.ac.uk/projects/fastqc/), trim the reads using TrimGalore! (https://www.bioinformatics.babraham.ac.uk/projects/trim_galore/) and align the reads using bowtie2-2.2.8 ^17^ with the following configurations: -X 1000 --local --fr --rg. Next, duplicate reads and insert size distributions were determined using picard-1.136. Additional quality metrics were generated using estimateReadsFiltering from deepTools ^18^ v3.0. MultiQC/1.3 ^19^ was then used to selectively combine the outputs from the above steps (Supplementary data 1-3).

### Filtering and Normalization

Coverage and normalization to the input was obtained using deepTools-2.5.4, bamCoverage (--normalizeTo1x 2451960000, --extendReads, --ignoreDuplicates, --blackListFileName) and bamCompare (--missingDataAsZero, --skipZeros, --blackListFileName) respectively. A mouse blacklist was obtained from the ENCODE project ^20^. A human blacklist was obtained from ENCODE (http://hgdownload.cse.ucsc.edu/goldenPath/hg19/encodeDCC/wgEncodeMapability/) and supplemented with unplaced contigs and decoy sequences from the 1000 genome projects. FRiPs scores were obtained using deepTools/plotEnrichment ^18^, the used peaks were taken from ENCODE projects (GSM1003519, GSM733645, GSM733685, GSM733737, GSM733743, GSM733754, GSM798321, GSM803499, GSM822287, GSM935545). In all downstream analyses, duplicates and blacklisted regions were removed, and all results were computed using fragments (parameter--extendReads).

### Visualization

Genome track plots were generated using the normalized bigWig files returned from bamCompare using deepTools/pyGenomeTracks (http://github.com/deeptools/pyGenomeTracks). All heatmaps and profiles were generated using deepTools. Sharp marks (CTCF, p300, H3K4me3, H3K27ac) were aligned around the center of the peak (computeMatrix reference-point), while other marks were considered broad and scaled to the region (computeMatrix scale-regions). In each case, flanking regions were added to show the signal away from enriched regions. computeMatrix was executed with the following parameters: --missingDataAsZero, --skipZeros, and --blackListFileName, while flanking regions (-a,-b) were defined according to the nature of the mark (broad 50kb, narrow 3kb).

### PCA plots

Principal component analysis was performed on the log2 ratio of the ChIP to the input (bigWig). First, for each sample a genome wide coverage was computed in windows of 10kb (deepTools/multiBigwigSummary bins, bs = 10000) and blacklisted regions were filtered. In R/3.4.1, prcomp was used to compute the PCA over the top 5000 most variable bins.

### Identification of restriction site position

Restriction sites were identified using the Bio.Restriction package from Biopython on the human assembly GRCh37 and mouse assembly GRCm38. In total, 44,340,181 sites and 39,483,329 sites were identified, respectively.

### Computation of restriction site bias

To identify if restriction site frequencies varied at different genomic regions, we computed the overlap between the total set of restriction sites (RS) and the different chromatin states for HepG2 ^21^ using coverageBed from bedtools2/2.27.0 ^22^. The number of RS per kbp was then computed using the sum of all observed overlaps divided by the sum of the respective peak length in kbp.

### Computation of RPKM for ChIPs

Read enrichment in HepG2 was assessed via FRiP scores. First, using deeptools/plotEnrichment (--extenReads --ignoreDuplicates), the fraction of reads, that fall into peaks was computed and obtained as a percentage. The FRiP scores were divided by the fraction of the genome falling into peaks and then multiplied by 1M to obtain RPKM values that are comparable across regions.

### Peak calling

Sharp peaks were identified using MACS2/2.1.0 ^11^, with the following parameters: -f BAM, --nomodel, -g 2900338458. The parameter --extsize was given as the median fragment length computed using Picard. For broad marks the peaks were called using histoneHMM/histoneHMM_call_regions.R ^23^ with bin size = 750 and -P 0.1.

### Chromatin states

Chromatin states were obtained from the UCSC table browser ^24^. The 15 chromatin states were obtained using chromHMM for the HepG2 cell line ^21^. Chromatin state 15 was not used, as it mostly contained human blacklisted regions.

### Code availability

The basic mapping and QC was done using v0.5 of snakePipes ^25^ for DNA-mapping and ChIP-seq, with only a minor modification to adjust for local mapping in the Bowtie2 step (*--local*). The workflows are freely avaliable at https://github.com/maxplanck-ie/snakemake_workflows. The RELACS demultiplexing step was done via demultiplex_relacs.py/version 1.0, which is available at https://github.com/dpryan79/Misc/tree/master/InternalBarcodes. For customized downstream analysis we used deepTools ^18^ v 2.5.4, as described above.

